# NECAPs are negative regulators of the AP2 clathrin adaptor complex

**DOI:** 10.1101/192898

**Authors:** Gwendolyn M Beacham, Edward A Partlow, Jeffrey J Lange, Gunther Hollopeter

## Abstract

Eukaryotic cells internalize transmembrane receptors via clathrin-mediated endocytosis, but it remains unclear how the machinery underpinning this process is regulated. We recently discovered that membrane-associated muniscin proteins such as FCHo and SGIP initiate endocytosis by converting the AP2 clathrin adaptor complex to an open, active conformation that is then phosphorylated (Hollopeter et al., 2014). Here we report that loss of *ncap-1*, the sole *C*. *elegans* gene encoding an adaptiN Ear-binding Coat-Associated Protein (NECAP), bypasses the requirement for FCHO-1. Biochemical analyses reveal AP2 accumulates in an open, phosphorylated state in *ncap-1* mutant worms, suggesting NECAPs promote the closed, inactive conformation of AP2. Consistent with this model, NECAPs preferentially bind open and phosphorylated forms of AP2 *in vitro* and localize with constitutively open AP2 mutants *in vivo*. NECAPs do not associate with phosphorylation-defective AP2 mutants, implying that phosphorylation precedes NECAP recruitment. We propose NECAPs function late in endocytosis to inactivate AP2.

## INTRODUCTION

Eukaryotic cells internalize transmembrane protein cargo, such as laden receptors, by enrobing cargo-containing regions of the plasma membrane with a cytosolic clathrin coat. The GTPase dynamin releases the nascent transport vesicle into the cytosol for subsequent delivery to internal organelles. This fundamental cellular process is called clathrin-mediated endocytosis. The Adaptor Protein 2 (AP2) complex, a heterotetramer of *α*, *β*2, μ2, and *σ*2 subunits (Figure 1– figure supplement 1A), actively couples polymerization of the clathrin coat to membrane phospholipids and cargo molecules destined for internalization (Kirchhausen, Owen, & Harrison, 2014). In this manner, AP2 choreographs turnover of the cell surface and entry into the endolysosomal system, yet it remains unclear how this complex is regulated with spatiotemporal precision to maintain appropriate levels of endocytosis.

AP2 activity is likely regulated at the structural level. Biochemical (Matsui & Kirchhausen, 1990; Rapoport et al., 1997) and structural data suggest that AP2 adopts at least two functionally distinct conformations (Figure 1–figure supplement 1B). In one arrangement, the binding pockets for membrane, cargo, and clathrin are occluded; this orientation is thought to represent a closed, inactive state (Collins, McCoy, Kent, Evans, & Owen, 2002). Molecular rearrangement of the AP2 subunits exposes these binding sites and results in an open, active complex that presumably coordinates the formation of endocytic pits (Jackson et al., 2010; Kelly et al., 2014; Kelly et al., 2008).

Multiple inputs coordinate the conformational rearrangement of AP2 but recent data suggest that membrane-associated muniscin proteins (Reider et al., 2009) allosterically activate the AP2 complex (Hollopeter et al., 2014; Umasankar et al., 2014). These include SH3-containing GRB2-like protein 3-Interacting Protein (SGIP) (Uezu et al., 2007) and Fer/CIP4 Homology domain only (FCHo) proteins (Henne et al., 2010; Sakaushi et al., 2007). In *C. elegans*, loss of FCHO-1 causes AP2 to dwell in the inactive state, leading to hindered endocytosis (Hollopeter et al., 2014). Gain-of-function mutations in AP2 that selectively destabilize the closed conformation, thereby artificially inducing the open state, can bypass loss of FCHO-1. A conserved region of muniscins called the AP2 activator domain (APA), binds AP2 and is sufficient to rescue *fcho-1* mutants (Hollopeter et al., 2014) and FCHo-deficient tissue culture cells (Umasankar et al., 2014).

Activation of AP2 and subsequent clathrin binding also require additional molecular events, including interactions with PI(4,5)P_2_ and cargo (Ehrlich et al., 2004; Jackson et al., 2010; Kadlecova et al., 2017; Kelly et al., 2014; Kelly et al., 2008). Phosphorylation of the AP2 μ2 subunit by an AP2-Associated Kinase (AAK1) (Conner & Schmid, 2002) is also proposed to be required for endocytosis (Olusanya, Andrews, Swedlow, & Smythe, 2001; Ricotta, Conner, Schmid, von Figura, & Honing, 2002) and is associated with the open form of AP2 (Hollopeter et al., 2014; Honing et al., 2005). However, it is not entirely clear whether phosphorylation induces the open state, stabilizes the open state (Jackson et al., 2010; Kadlecova et al., 2017), or marks adaptor complexes that have already been incorporated into clathrin coats (Conner, Schroter, & Schmid, 2003; Jackson et al., 2003; Semerdjieva et al., 2008).

How then is AP2 inactivated and returned to the cytosol after endocytosis is complete? The heat shock protein Hsc70 (Chappell et al., 1986) and its cofactors (Greener, Zhao, Nojima, Eisenberg, & Greene, 2000; Umeda, Meyerholz, & Ungewickell, 2000; Ungewickell et al., 1995) remove clathrin coats from vesicles. Release of AP2 is also thought to depend on Hsc70 (Hannan, Newmyer, & Schmid, 1998) but appears to require additional factors that stimulate the dephosphorylation of μ2 (Ghosh & Kornfeld, 2003) and the conversion of vesicular PI(4,5)P_2_ into PI(4)P (Cremona et al., 1999; Semerdjieva et al., 2008). Whether there are mechanisms to directly restore the inactive, closed conformation of AP2 has been relatively unexplored.

In this study, we identify adaptiN Ear-binding Coat-Associated Proteins (NECAPs) as AP2 modulators that promote inactivation of the complex. NECAPs were originally discovered through proteomic analysis of clathrin-coated vesicles and were shown to bind the AP2 α appendage (Ritter et al., 2003). Using an unbiased genetic screen in *C*. *elegans*, we have discovered that NECAPs work in opposition of muniscins. We demonstrate that in worms lacking NECAPs (*ncap-1* mutants), AP2 accumulates in an open, hyper-phosphorylated state and that heterologous NECAPs restore the closed conformation. NECAPs bind open and phosphorylated forms of the AP2 core but not phosphorylation-defective AP2 mutants, suggesting that NECAPs target the active complex. Together, our genetic and biochemical evidence establish NECAPs as negative regulators of AP2.

## RESULTS

### Loss of *ncap-1* suppresses *fcho-1* mutants

By mutagenizing worms lacking *fcho-1* and selecting for offspring that outcompete their siblings (Hollopeter et al., 2014), we isolated ten independent loss-of-function mutations in *ncap-1,* which encodes the sole adaptiN Ear-binding Coat-Associated Protein (NECAP) in *C*. *elegans* (Figure 1A). Worms with null mutations in *fcho-1* exhibit reduced fitness and require twice as long as wild type worms to populate a culture plate and consume the bacterial food source. Additionally, they display a distinctive “jowls” phenotype that is indicative of compromised AP2 activity (Gu et al., 2013; Hollopeter et al., 2014). Loss of NCAP-1 suppressed the jowls phenotype (Figure 1B) and ameliorated the fitness of *fcho-1* mutants (Figure 1C). Expression of fluorescently-tagged NCAP-1 in *fcho-1 ncap-1* double mutants restored the *fcho-1* phenotype (Figure 1B and C), confirming that suppression of *fcho-1* is due to loss of NCAP-1 function.

**Figure 1.**
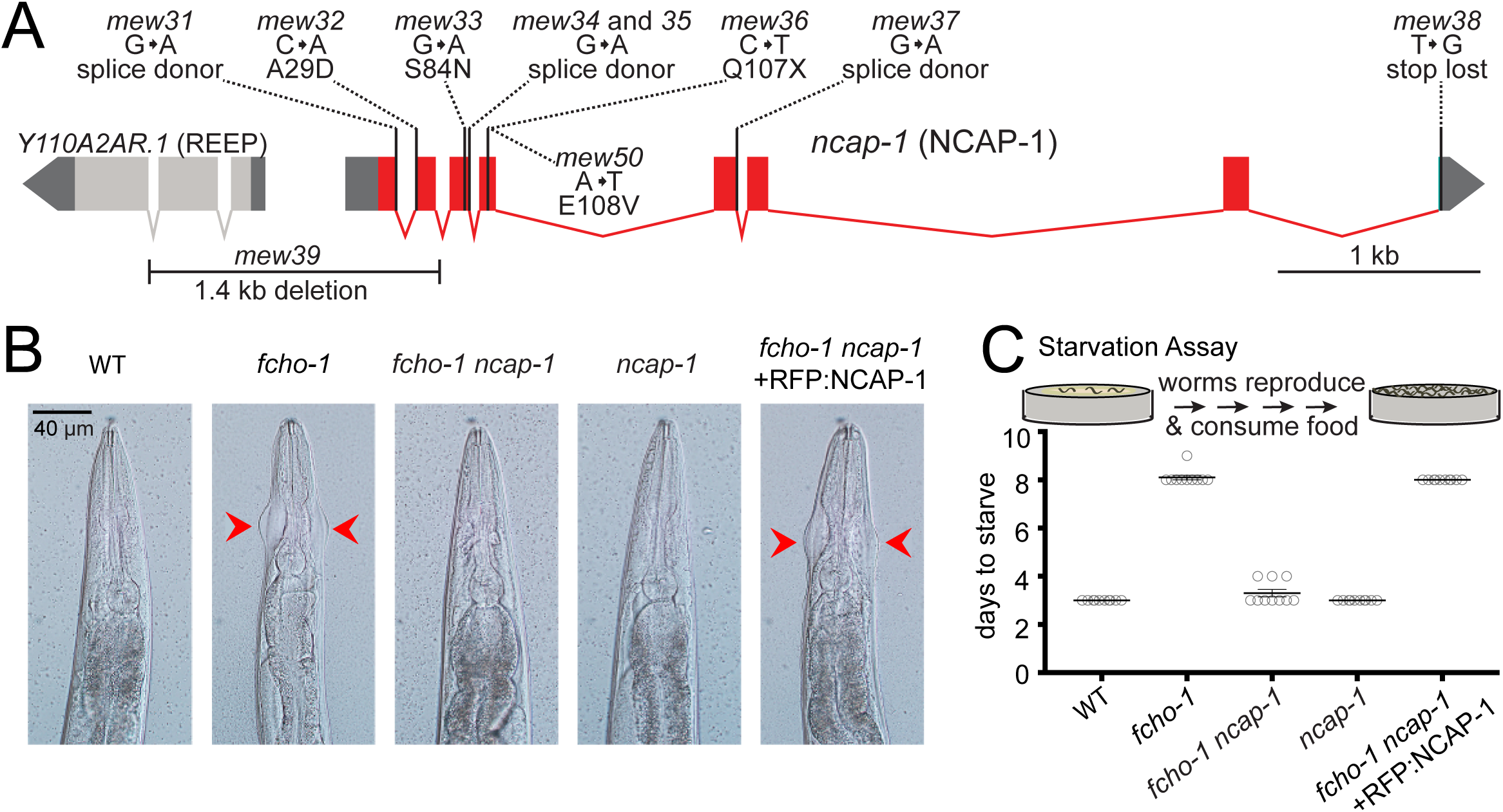
Loss of NCAP-1 suppresses *fcho-1* mutants. **A.** Gene model of the *C. elegans ncap-1* locus. Boxes represent exons. Mutations isolated from the *fcho-1* suppressor screen are indicated. The deletion allele, *mew39,* was used throughout this study as *ncap-1*. The neighboring gene (Y110A2AR.1) is predicted to encode a receptor expression-enhancing protein (REEP). **B.** Representative images of worms. Red arrowheads indicate jowls phenotype. Anterior is up. Wild type (WT); red fluorescent protein-tagged NCAP-1 (RFP:NCAP-1). **C.** Starvation assay. Data (bottom) represent days required for worms to reproduce and consume bacterial food source (top schematic). Bars indicate mean ± SEM for n = 10 biological replicates.

Previously we observed that suppression of *fcho-1* correlates with recovery of AP2 activity (Hollopeter et al., 2014). To evaluate if loss of NCAP-1 also increases AP2 activity in *fcho-1* mutants, we imaged GFP-tagged AP2 α adaptin (APA-2:GFP) in macrophagic cells called coelomocytes that exhibit robust levels of endocytosis (Sato, Norris, Sato, & Grant, 2014). Muniscins, such as FCHO-1, stabilize AP2 on the plasma membrane to promote its incorporation into presumptive pits, or clusters (Cocucci, Aguet, Boulant, & Kirchhausen, 2012; Henne et al., 2010; Hollopeter et al., 2014). We performed Fluorescence Recovery After Photobleaching (FRAP) to quantify AP2 stability on the coelomocyte membrane. In the *fcho-1* mutants, fluorescent AP2 recovers approximately three times faster than in wild type worms, indicating that AP2 association with the membrane is destabilized (Hollopeter et al., 2014). Loss of NCAP-1 slowed AP2 kinetics in *fcho-1* animals (Figure 2A), suggesting that AP2 is incorporated into longer-lived structures. Indeed, we observed improved AP2 clustering in some cells, but the trend was not significant (Figure 2B). Endocytosis of an AP2-dependent model cargo is compromised in *fcho-1* mutants (Hollopeter et al., 2014) whereas cargo internalization was partially rescued in *fcho-1 ncap-1* worms (Figure 2C). These data indicate that a loss of NCAP-1 improves AP2 activity in *fcho-1* mutants.

**Figure 2.**
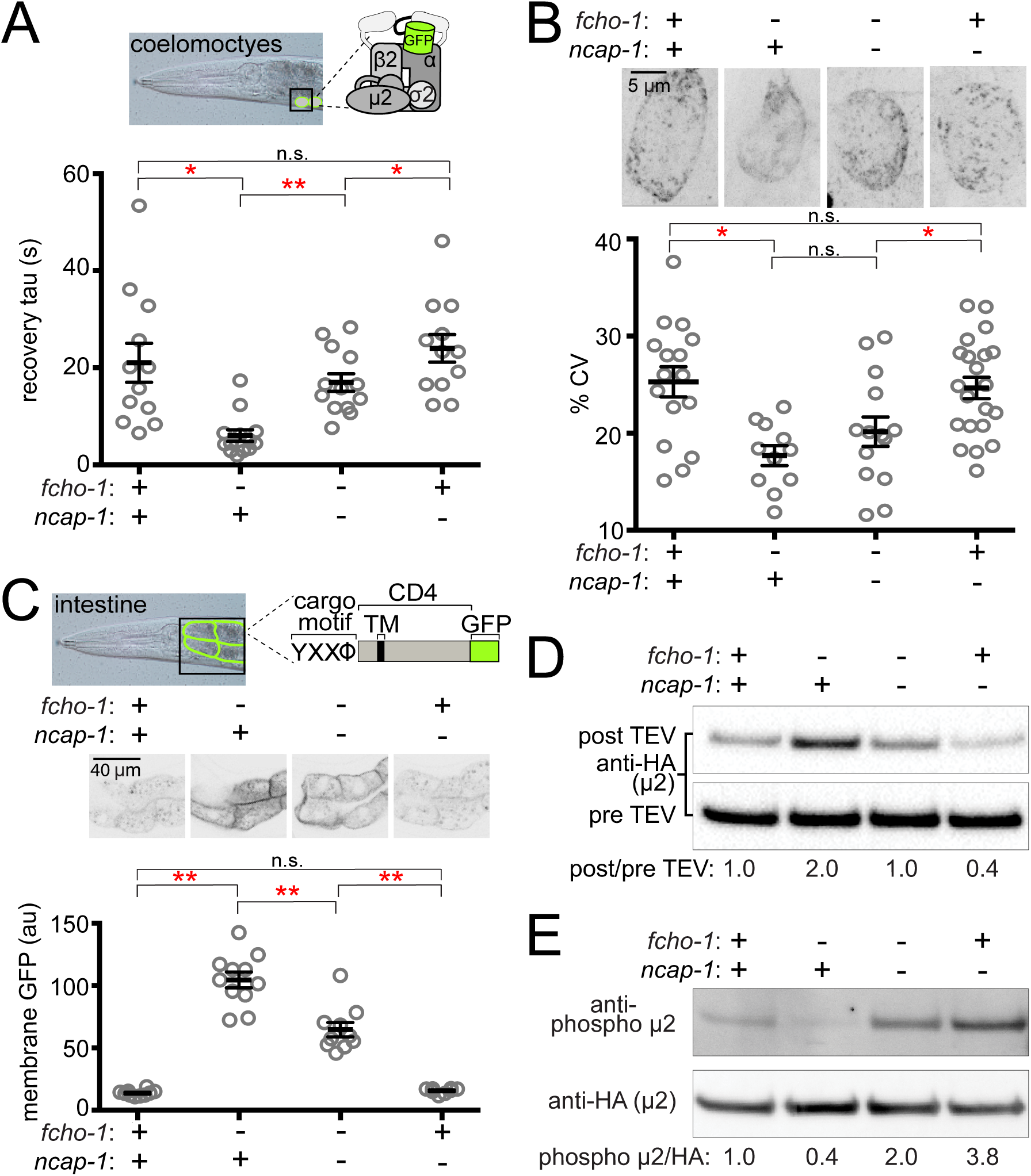
Loss of NCAP-1 restores AP2 activity in *fcho-1* mutants. **A.** FRAP analysis of GFP-tagged AP2 α adaptin (APA-2:GFP) on membranes of coelomocytes (top schematic). Time constants (tau) of the fluorescence recovery are plotted (below). **B.** AP2 localization in coelomocytes. Representative confocal images of coelomocytes in worms expressing APA-2:GFP. Micrographs (top) are representative maximum projections of Z-slices through approximately half of a cell. Data (bottom) represent the coefficient of variance (%CV) of pixel intensities for individual cells. **C.** Cargo assay. Representative confocal micrographs of intestinal cells (middle) in worms expressing a P-tagged artificial AP2 cargo (top schematic). The average pixel intensity along a basolateral membrane was measured (bottom). **A-C.** Bars indicate mean ± SEM for n ≥ 8 biological replicates. * p<0.05, ** p<0.001, not nificant (n.s.), unpaired, two-tailed T-test. **D.** Protease-sensitivity assay. Western blot analysis of whole worm lysates was used to quantify the amount of full-length μ2 (anti-HA, 50 kDa) before (pre TEV, bottom blot) and after protease induction (post TEV, top blot). Band intensities were compared to a tubulin loading control and normalized to *fcho-1*(+) *ncap-1*(+) ratio (values below). **E.** Phosphorylation assay. Western blot analysis of whole worm lysates quantify phosphorylated μ2 (top blot) relative to total μ2 subunit (bottom blot). Values indicate band intensity ratios of phospho μ2 compared to total μ2, normalized to the *fcho-1*(+) *ncap-1*(+) ratio (values below). **D and E.** Blots are resentative examples of ≥ 3 biological replicates. +, wild type allele; -, deletion allele.

### Open and phosphorylated AP2 accumulates in *ncap-1* mutants

We previously discovered that AP2 activity in *fcho-1* mutants is also partially restored by amino acid substitutions that specifically destabilize the closed conformation of the adaptor complex, henceforth referred to as “open AP2 mutations” (Hollopeter et al., 2014). Because loss of NCAP-1 phenotypically mimicked open AP2 mutations, we tested whether AP2 also dwells in the open state in *fcho-1 ncap-1* worms. We evaluated AP2 conformation using an *in vivo* protease sensitivity assay (Hollopeter et al., 2014). Briefly, a Tobacco Etch Virus (TEV) protease site was inserted into a surface loop of the μ2 subunit that becomes more exposed when AP2 is opened (Jackson et al., 2010; Matsui & Kirchhausen, 1990). Following induction of TEV protease expression, the ensemble protease-sensitivity of the AP2 complexes is determined using western blot analysis of μ2 in whole worm lysates. In *fcho-1* worms, AP2 dwells in a protease-insensitive, closed state (Hollopeter et al., 2014) and AP2 was more protease-sensitive (open) in *ncap-1* mutants (Figure 2D). These results explain the phenotypic rescue of *fcho-1* mutants and indicate that AP2 accumulates in an open state in the absence of NCAP-1.

Phosphorylation of μ2 correlates with open AP2 in *C. elegans* (Hollopeter et al., 2014). Because loss of NCAP-1 increases AP2 protease-sensitivity, we wondered whether μ2 phosphorylation might also be elevated. We quantified the phosphorylation status of AP2 *in vivo* by blotting worm lysates with an antibody specific to the phosphorylated threonine in μ2 (T160). Loss of NCAP-1 resulted in T160 phosphorylation levels greater than in wild type worms (Figure 2E). Thus, AP2 adopts a hyper-phosphorylated, open state without the action of NCAP-1.

### Negative regulation of AP2 is a conserved function of NECAPs

While most invertebrate and fungal genomes encode a single NECAP, vertebrates express two closely-related forms (Dergai, Iershov, Novokhatska, Pankivskyi, & Rynditch, 2016; Manna, Gadelha, Puttick, & Field, 2015) that are thought to be functionally distinct. Brain-enriched NECAP1 is proposed to modulate endocytic pit formation by regulating the binding of clathrin to the AP2 β2 linker and the coordination of accessory protein recruitment to the α appendage (Ritter et al., 2013). By contrast, ubiquitously-expressed NECAP2 has been proposed to recruit a different clathrin adaptor, AP1, to early endosomes in order to facilitate fast recycling of receptors back to the cell surface (Chamberland, Antonow, Dias Santos, & Ritter, 2016). To determine if NECAPs might share the ability to inactivate AP2 in the context of our *C*. *elegans* system, we expressed heterologous NECAPs as single-copy transgenes in *fcho-1 ncap-1* mutant worms. Mouse NECAP1 and NECAP2, as well as a NECAP from the multicellular fungus, *Sphaerobolus stellatus,* recapitulated the reduced fitness of *fcho-1* mutants (Figure 3A) and caused AP2 to adopt a more closed, protease-insensitive state (Figure 3B). It appears that NECAPs have retained the ability to negatively regulate AP2, at least in *C. elegans*.

**Figure 3.**
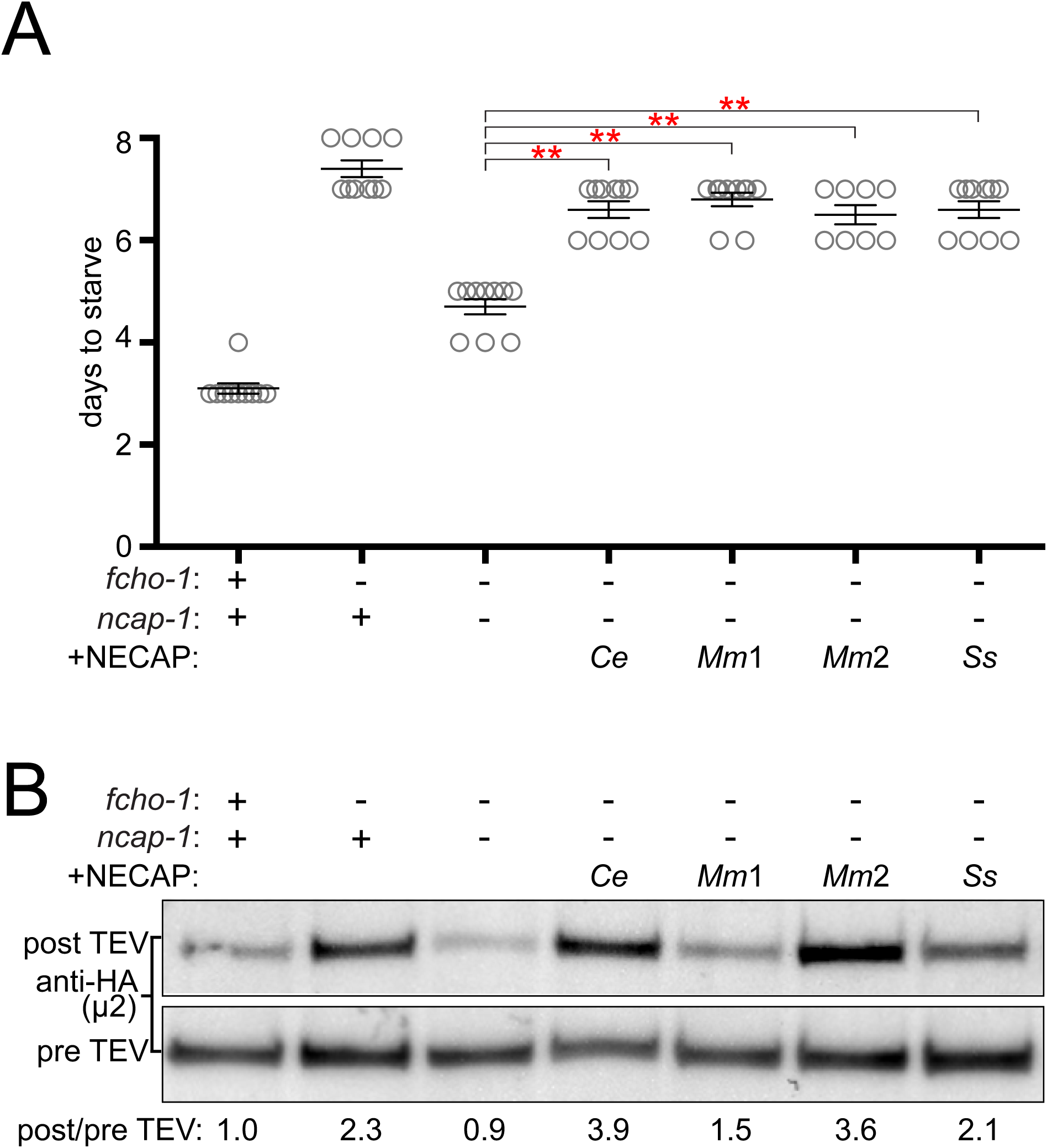
NECAPs restore closed AP2 in *fcho-1 ncap-1* worms. NECAPs were expressed as single copy transgenes in *fcho-1 ncap-1* worms. *Ce*, *C. elegans*; *Mm*, *M. musculus*; *Ss*, *Sphaerobolus stellatus* (multicellular fungus). +, wild type allele; -, deletion allele. **A.** Starvation assay performed as in Figure 1C. Bars represent mean ± SEM for n ≥ 7 biological replicates. ** p<0.001, unpaired, two-tailed T-test. **B.** Protease-sensitivity assay as in Figure 2D. Blot is representative of 2 biological replicates.

### NECAPs bind the open and phosphorylated AP2 core *in vitro*

We hypothesized that NECAPs might negatively regulate AP2 by binding directly to the complex. Indeed, affinity-tagged NECAPs co-purified endogenous AP2 complexes from HEK293 cells (Figure 4A), consistent with previous reports (Ritter et al., 2003). NECAPs are thought to bind the AP2 α adaptin appendage domain and the clathrin binding box in the β2 adaptin hinge region via a C-terminal WXXF motif and an N-terminal PHear domain, respectively (Ritter et al., 2013; Ritter et al., 2003). However, we were interested to examine whether NECAPs also bind to the AP2 core (Figure 1–figure supplement 1B) because this would offer a direct route to conformational regulation. We purified recombinant vertebrate AP2 cores lacking ears and linkers and tested binding of these cores to recombinant NECAPs from worms and mice (Figure 4B). Interestingly, NECAPs did not bind unmodified AP2 cores (Figure 4B).

**Figure 4.**
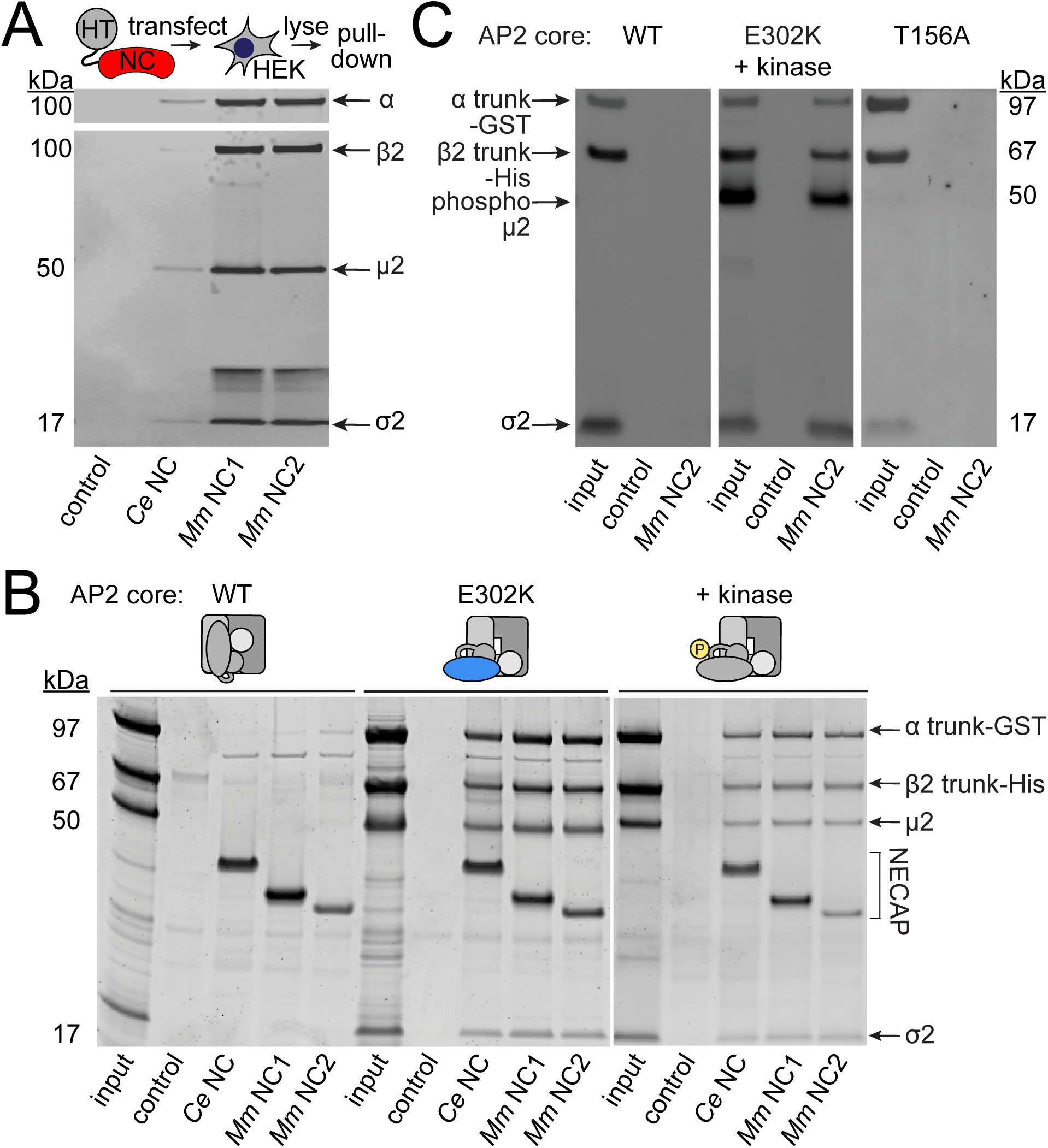
NECAPs bind the open and phosphorylated AP2 core. Pulldown assays using affinity-tagged NECAPs. Proteins were cleaved from the affinity tag (HaloTag), electrophoretically separated and then blotted for AP2 subunits (A and C) or stained prior to imaging (B). Control, HaloTag alone; NC, NECAP; *Ce, C. elegans*; *Mm*, *M. musculus*. **A.** Western blot analysis (middle) of samples purified from human cell lysates (top schematic) expressing the indicated heterologous NECAP bait (bottom). **B and C.** *In vitro* pulldown assays using purified recombinant bait (NECAPs, bottom) and prey (vertebrate AP2 cores, top). Co-expression with the kinase domain from mouse AAK1 (+ kinase) generates phosphorylated AP2. Amino acid changes in μ2 are indicated: E302K, constitutively open AP2; T156A, phosphorylation-defective AP2; see also Figure 1–figure supplement 1C. **A-C.** Data are representative of 2 biological (A), 1 technical (B), and 2 technical (C) replicates.

Because open and phosphorylated AP2s accumulate in *ncap-1* mutants, we reasoned that NECAPs might act upon these modified forms of AP2. To test our hypothesis, we introduced a previously-characterized open AP2 mutation in the μ2 subunit (E302K; Figure 1– figure supplement 1C) (Hollopeter et al., 2014) to produce recombinant AP2 cores that dwell in the open state. We also co-expressed AP2 with the kinase domain of AAK1 to purify AP2 cores with phosphorylated μ2(T156) (Honing et al., 2005). When these modified complexes were tested in our pulldown assays, we observed that NECAPs bound both open and phosphorylated AP2 (Figure 4B and C). These data suggest that NECAPs associate with the AP2 complex in a conformation-dependent manner.

### NCAP-1 localizes with constitutively open AP2 *in vivo*

To corroborate our *in vitro* pulldown results we sought *in vivo* evidence that NECAPs associate with open and phosphorylated AP2. In *C. elegans*, fluorescently-tagged AP2 is enriched at the nerve ring, a major neuropil of bundled axons encircling the pharynx (White, Southgate, Thomson, & Brenner, 1986) (Figure 5). We used confocal microscopy to observe localization of NCAP-1 tagged with a red fluorescent protein (RFP:NCAP-1) at the nerve ring in live worms. In wild type worms, NCAP-1 was not overtly enriched compared to AP2. However, introduction of a mutation in μ2 known to generate hyper-phosphorylated, constitutively open AP2 (E306K; Figure 1–figure supplement 1C) (Hollopeter et al., 2014) enhanced localization of NCAP-1 nearly 2-fold (Figure 5). These data suggest that NECAP favors association with activated forms of AP2 *in vivo*.

**Figure 5.**
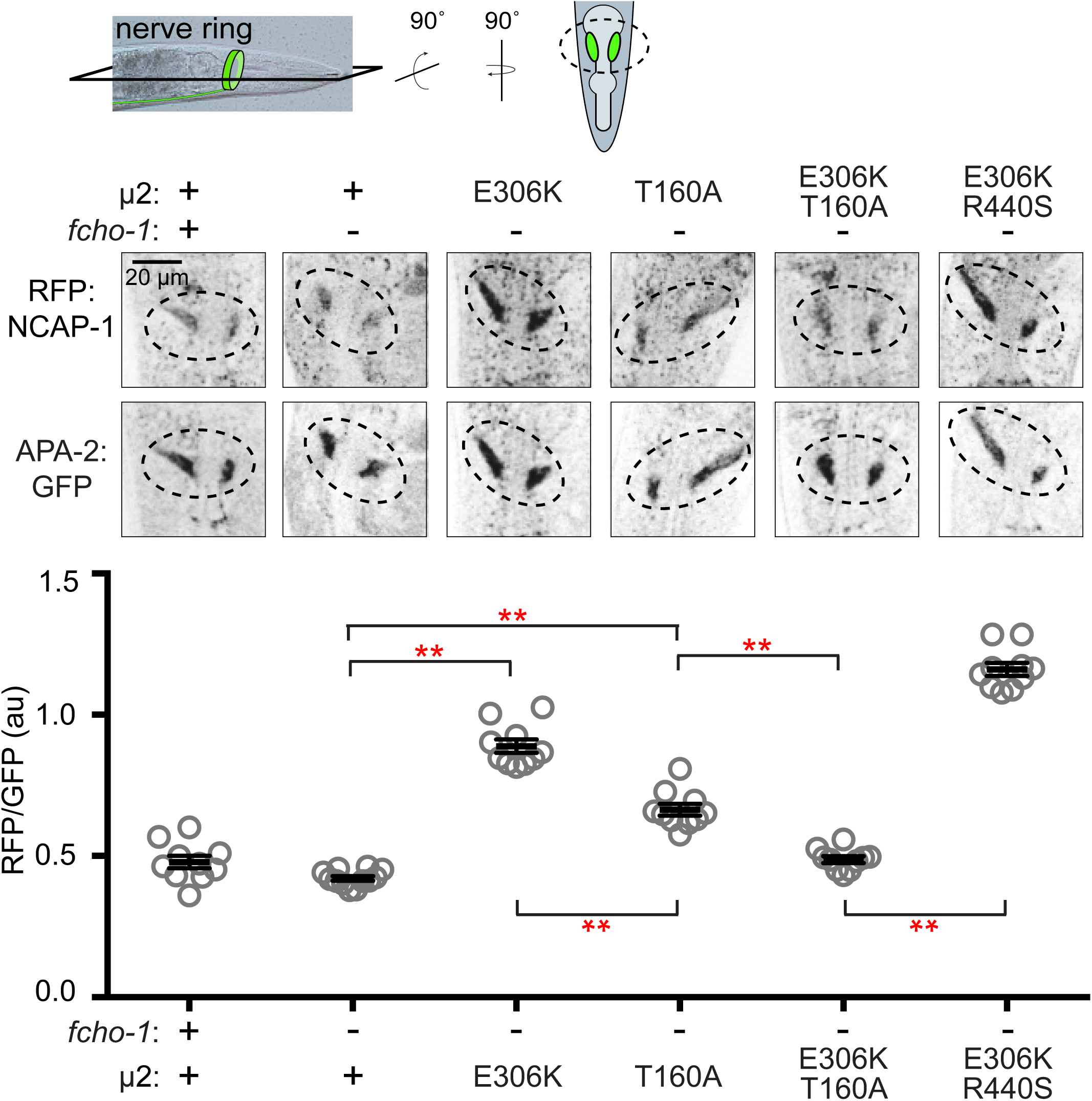
Phosphorylated AP2 recruits NCAP-1 *in vivo*. Representative confocal slices (middle) through the approximate center of the nerve ring of worms (top schematic) expressing RFP:NCAP-1 and APA-2:GFP. RFP to GFP signal intensity at the nerve ring is plotted (bottom). Mutations in μ2 are indicated: E306K and R440S, constitutively open AP2; T160A, phosphorylation-defective AP2; see also Figure 1–figure supplement 1C. Bars indicate mean ± SEM for n ≥ 10 biological replicates. ** p<0.001, unpaired, two-tailed T-test. +, wild type allele; -, deletion allele.

### AP2 phosphorylation site mutations weaken NECAP binding

We were curious whether phosphorylation of AP2 is a prerequisite for NECAP binding. Interestingly, in *C*. *elegans*, mutating the phosphorylation site, μ2(T160), to an alanine, isoleucine, proline, or glutamate suppresses *fcho-1* and produces an open complex according to the *in vivo* protease-sensitivity assay (Hollopeter et al., 2014). However, when we purified vertebrate AP2 cores containing the phosphorylation site mutation (T156A; Figure 1–figure supplement 1C) and tested their ability to bind mouse NECAP2, we observed very little interaction in pulldown assays (Figure 4C). This result suggests that NECAPs do not bind phosphorylation-defective AP2. To determine whether the phosphorylation site mutants also impact NCAP-1 association with AP2 *in vivo*, we examined the localization of RFP:NCAP-1 in μ2(T160A) mutant worms. Compared to mutants with hyper-phosphorylated AP2 (E306K), the T160A mutants had less NCAP-1 at the nerve ring (Figure 5). These results are consistent with the model that NCAP-1 associates with AP2 in a phosphorylation-dependent manner.

The preferential association of NCAP-1 with the hyper-phosphorylated AP2 (E306K) compared to the phosphorylation-defective AP2 (T160A) was not simply due to differences in the extent to which these mutations generate a protease-sensitive, open AP2 complex (Hollopeter et al., 2014). When the E306K and T160A mutations were both present in the same μ2 subunit, less RFP:NCAP-1 was localized to the nerve ring compared to when the E306K mutation was combined with another open AP2 mutation (R440S; Figure1–figure supplement 1C and Figure 5) (Hollopeter et al., 2014). In other words, the effect of the phosphorylation site mutation was dominant, while the two open AP2 mutations were additive, with respect to NCAP-1 association. Thus, phosphorylation site mutations preclude the association of NCAP-1 with open AP2.

## DISCUSSION

In *C*. *elegans*, loss-of-function mutations in *ncap-1* bypass the requirement for FCHO-1. These results could be consistent with two distinct models: 1) NCAP-1 maintains the closed conformation of AP2 until FCHO-1 releases this inhibition, or 2) NCAP-1 generates the closed conformation of AP2, either directly or indirectly, and this form predominates in the absence of FCHO-1. The results of our biochemical and imaging data are consistent with the latter; NECAPs bind the open, phosphorylated AP2 core and restore the closed form of the complex *in vivo*. We propose that NECAPs act downstream of muniscin function and μ2 phosphorylation to inactivate AP2 (Figure 6).

**Figure 6.**
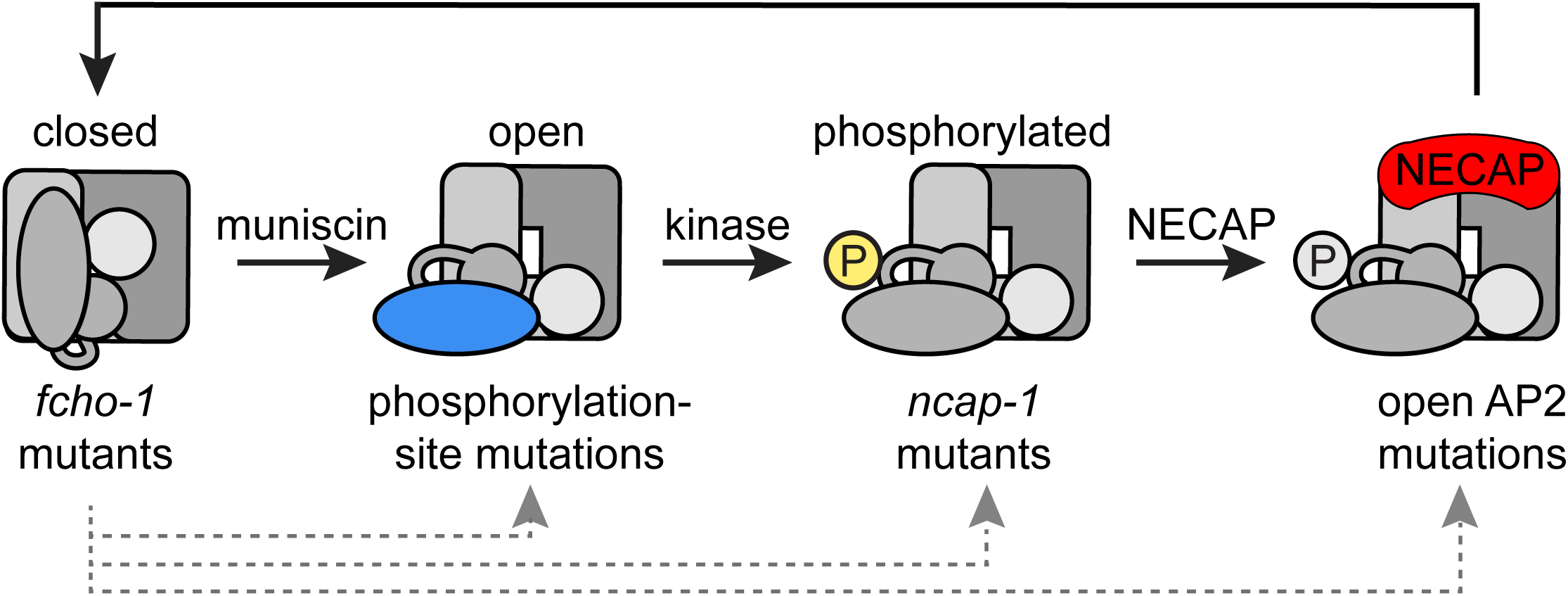
Model of AP2 activation and inactivation. Three classes of mutations increase AP2 activity in the absence of FCHO-1 (bottom), promoting accumulation of the complex at the indicated steps in the cycle.

### *fcho-1* suppressors disrupt the AP2 inactivation pathway

In the absence of FCHO-1, nematodes exhibit reduced fitness and AP2 dwells in a closed, hypo-phosphorylated state. Using an unbiased genetic screen, we isolated mutations that independently improve the fitness of *fcho-1* genetic nulls. These *fcho-1* suppressors occur in three distinct classes: dominant missense mutations that (1) disrupt the closed conformation of AP2 or (2) mutate the phosphorylation site on μ2, and (3) recessive mutations that inactivate NCAP-1. We propose that these mutations rescue *fcho-1* by restoring AP2 activity, but that they achieve this through different mechanisms. Previously, the phosphorylation site mutants appeared to behave similarly to open AP2 mutants in our assays (Hollopeter et al., 2014). However, we now know NECAPs distinguish between open and phosphorylation-defective AP2. Open, hyper-phosphorylated AP2 mutants bind NECAPs but resist inactivation. By contrast, phosphorylation site mutants are likely resistant to inactivation because they evade NECAP binding. Importantly, our data suggest that the suppressors all bypass FCHO-1 by disrupting an inactivation process by which the closed conformation of AP2 is normally restored. The mutations also appear to cause the adaptor complex to accumulate in active forms at discrete steps along this recycling pathway (Figure 6).

It is curious that the unnaturally active forms of AP2 isolated from our *fcho-1* suppressor screen do not exhibit enhanced endocytosis or membrane association when the *fcho-1* gene is intact (Hollopeter et al., 2014). In other words, why is there not a functional consequence associated with hyperactive AP2 in an otherwise wild type background? Perhaps there are compensatory mechanisms that prevent rampant endocytosis or, alternatively, our assays may lack the sensitivity and range necessary to detect the differences. Indeed, another group has reported that endocytic pits become enlarged after knockdown of NECAP1 and these structures could represent a consequence of overactive AP2 (Ritter et al., 2013).

### Conformation-dependent regulation of AP2

NECAPs form stable complexes *in vitro* with open and phosphorylated AP2 cores, suggesting that activated AP2 may be the endogenous substrate of NECAPs. In vertebrates, NECAP1 has been reported to bind the AP2 α ear and the β2 linker early in endocytosis to regulate the recruitment of endocytic accessory proteins and to compete for clathrin binding, respectively (Ritter et al., 2004; Ritter et al., 2013). It is unclear whether these events are concurrent with the conformation-dependent binding of NECAPs to the AP2 core. Because the clathrin binding site is thought to be exposed when AP2 binds membranes (Kelly et al., 2014) it is possible NECAPs could inhibit clathrin binding upon association with active AP2 (Ritter et al., 2013). The AP2 appendages can exhibit non-specific binding *in vitro* (Hollopeter et al., 2014) so it has been challenging to evaluate this model using purified components. However, imaging data suggest NECAP dynamics during endocytic pit formation mimic those of clathrin (Taylor, Perrais, & Merrifield, 2011), consistent with our model that AP2 activation precedes NECAP recruitment.

The accumulation of phosphorylated AP2 in the absence of NCAP-1 indicates thatphosphorylation also precedes NECAP activity but is it unclear whether the phospho-threonine is an essential feature of the NECAP-AP2 interface. Changing the threonine to aphosphorylation-defective alanine does appear to disrupt the association of NECAPs with AP2. However, we have not excluded the possibility that this mutation simply precludes NECAP binding by altering the conformation of AP2. Indeed, we observed that NECAPs bind open AP2 cores that have not been phosphorylated (the E302K cores, Figure 4B), further supporting the model that NECAPs bind to a face of active AP2 that does not include the phosphorylated threonine.

### The core function of NECAPs

While NECAPs are widely conserved across eukaryotic organisms (Dergai et al., 2016), the precise function of this protein family has remained enigmatic. Vertebrate NECAPs exhibit different tissue distributions and are proposed to function in mutually exclusive pathways. For example, NECAP2 was recently shown to regulate AP1 instead of AP2 (Chamberland et al., 2016; Ritter et al., 2013) Despite this proposed divergence, both vertebrate NECAPs rescue loss of NCAP-1 in the nematode, as does a fungal NECAP. These results indicate that the capacity to negatively regulate AP2 has been conserved among NECAPs.

How do NECAPs sustain the AP2 cycle? We propose the following model (Figure 6): muniscins such as FCHO-1 promote the open state of AP2, which is phosphorylated on the μ2 subunit by the AP2-associated kinase. NECAPs then bind the AP2 core to counteract the open state, presumably by facilitating a conformational change or dephosphorylation event later in endocytosis. This recycles AP2 back to its inactive state in the cytosol and renders the complex available for another round of endocytosis.

## MATERIALS AND METHODS

### Strains

*C. elegans* were maintained using standard procedures (Brenner, 1974) on nematode growth medium (NGM) plates seeded with *E. coli* (OP50). For a complete list of strains, see Supplementary file 1A.

### Data analysis

Unpaired, parametric, two-tailed T-tests were performed using GraphPad Prism (version 7.0c for Mac, GraphPad Software, La Jolla, CA, USA, http://www.graphpad.com).

### Identification of *ncap-1* mutants

Our initial mutagenesis of *fcho-1* null animals (Hollopeter et al., 2014) yielded four recessive suppressors in the same complementation group. Single nucleotide polymorphism mapping (Davis et al., 2005) placed them on chromosome II and whole genome sequencing revealed four independent mutations in the *ncap-1* gene (*mew31*, *mew36*, *mew38*, and *mew39*). Additional alleles were identified among subsequent *fcho-1* suppressors by amplifying and sequencing the coding segments of *ncap-1* with primer pairs oGH678-9 and oGH680-1. Strains and oligonucleotides are listed in Supplementary file 1.

### Starvation assay

The starvation assay was performed as previously described (Hollopeter et al., 2014) except that the assay was performed at 25°C for Figure 1C.

### Preparation of worms for microscopy

For confocal fluorescence microscopy, worms were mounted on 8-10% agarose pads in 3 μL of a 1:1 mix of a 1 μm polystyrene bead slurry (Polysciences, Warrington, PA) and 2X PBS pH 7.4 (Kim, Sun, Gabel, & Fang-Yen, 2013). For differential interference contrast microscopy (Figure 1B), worms were mounted on 5% agarose pads in PBS pH 7.4 with 20 mM sodium azide.

### AP2 localization and FRAP analysis in coelomocytes

Worms expressing APA-2:GFP (*oxSi254*) were imaged as previously described (Hollopeter et al., 2014). Strains are listed in Supplementary file 1A.

### Cargo assay

The cargo assay was performed essentially as previously described (Hollopeter et al., 2014). Worms expressing an artificial cargo (*oxSi484*) were imaged on a Ziess LSM 880 confocal microscope (Biotechnology Resource Center, Cornell University, Ithaca, NY) with a 40x water immersion objective. All strains were imaged in one session with the same laser settings. Images were analyzed in Fiji (Schindelin et al., 2012). A user defined line was drawn along the membrane between intestinal segments 2 and 3, or segments 3 and 4, and the average pixel intensity was measured along the line. Strains are listed in Supplementary file 1A.

### TEV protease-sensitivity assay

The TEV assay was performed essentially as previously described (Hollopeter et al., 2014). Post TEV samples were collected at 5-6 h following heatshock (34°C, 1 h). Each sample represents 100 L4 hermaphrodites lysed in 1X Bolt LDS Sample Buffer (Invitrogen, Carlsbad, CA) containing fresh dithiothreitol (DTT; ∼100 mM) by sonication (1 s pulses at 90-95% amplitude for 2-3 min) in a cup horn (Branson Ultrasonics Corporation, Danbury, CT) that was chilled to 4°C. Samples were heated for 10 min at 70°C prior to gel electrophoresis. All samples were re-sonicated following the 70°C denaturation step if any exhibited excessive viscosity. Strains are listed in Supplementary file 1A.

### Western blots and SYPRO staining

Precast polyacrylamide gels (Bolt 4-12% Bis-Tris, Invtrogen) were used for all SDS PAGE experiments. For western blot analysis, proteins were transferred from gels to PVDF Immobilon membranes (Merck Millipore, Tullagreen, Carrigtwohill, Co. Cork, Ireland) using the Pierce Power Blot Cassette system (Thermo Scientific, Rockford, IL). All blocking and antibody incubations (except anti-HA, see below) occurred in Odyssey Blocking Buffer (LI-COR, Lincoln, NE). Primary antibodies and dilutions included mouse anti-adaptin α (1:500, BD Biosciences, San Jose, CA, 610501), rabbit anti-AP2B1 (1:500, Abcam, Cambridge, MA, 151961), rabbit anti-AP2M1 phospho T156 (1:1000, Abcam 109397), rabbit anti-AP2S1 (1:4000, Abcam 128950), mouse anti-flag (1:1000, Sigma-Aldrich F3165), and mouse anti-tubulin (1:2000, Sigma-Aldrich T5168). Secondary antibodies included goat anti-mouse Alexa Fluor 488 (1:4000, Life Technologies, Eugene, OR, A11029), goat anti-rabbit Alexa Fluor 647 (1:2000, Life Technologies, A21244), StarBright Blue 700 goat anti-rabbit (1:5000, BioRad, Hercules, CA), and goat anti-mouse IRDye 800CW (1:20000, LI-COR, 925-32210).

Blots probed with the anti-HA-Horseradish peroxidase (HRP) antibody (1:500, Roche 12013819001, Mannheim, Germany), were blocked in Tris Buffered Saline + 0.01% Tween 20 (TBST) with 5% nonfat dry milk. The antibody was diluted in TBST with 1% milk and incubation occurred at room temperature for 1 h. SuperSignal West Dura Extended Duration Substrate (Thermo Scientific) was used to detect peroxidase.

Primary antibody incubations occurred for 1 h at room temperature or overnight at 4°C. Secondary antibody incubations occurred at room temperature for 30 min – 1 h in the dark.TBST was used for all washes. Blots were imaged using Bio-Rad ChemiDoc MP systems and band intensities were quantified using the associated ImageLab software.

We used SYPRO Ruby Protein Gel stain (Lonza, Rockland, ME) to visualize the *in vitro* pulldowns in Figure 4B. Gels were fixed in 50% methanol/7% acetic acid for 30 min and stained with SYPRO for 4-16 h at room temperature in the dark with gentle agitation. Gels were then washed 2 X 15 min in 10% methanol/7% acetic acid and then 2 X 5 min in water before imaging on a Bio-Rad ChemiDoc MP system.

### *C. elegans* NECAP transgenes

We generated *C. elegans* targeting vectors for expression of NECAPs as single copy-transgenes (Frokjaer-Jensen et al., 2008). For Figure 1 and 5, a mini-gene encoding NCAP-1 was constructed using the Multisite Gateway System (Invitrogen). The first 4 exons of the ncap-1 gene were amplified with primer pair oGH731+3, while the other half of the coding sequence was amplified from cDNA using primer pair oGH734+6. The two amplicons were recombined using the Gibson assembly reaction (Gibson et al., 2009) and then recombined with the [1-2] donor vector using BP clonase (Invitrogen). The entry clone was amplified with oGH698+738 and a worm codon-optimized TagRFP-T (Ed Boyden, MIT, Cambridge, MA) was amplified with oGH526+8 and inserted upstream of the *ncap-1* coding sequences using Gibson assembly. The resulting [1-2] entry clone (pGH505) was recombined with a [4-1] entry containing the ubiquitous dpy-30 promoter, the unc-54 3’UTR in a [2-3] entry and the [4-3] destination vector pCFJ150 (Christian Frøkjær-Jensen, University of Utah) using LR clonase (Invitrogen) to generate the MosSCI targeting vector (pGH495) that was injected into worm strain EG6699 (Frokjaer-Jensen, Davis, Ailion, & Jorgensen, 2012).

For Figure 3, the coding sequences of NECAPs were amplified from plasmid templates using the following primers: oEP366-7 for *C. elegans* NCAP-1 (NM_061997.5), oEP409-10 for *M. musculus* NECAP1 (BK000656.1) and oEP407-8 for *M. musculus* NECAP2 (BK000657.1).The coding sequence of NECAP from *Sphaerobolus stellatus* (Cannonball Fungus protein KIJ44287) was synthesized as a gBlock (IDT, Coralville, IA). These DNA fragments were then assembled in a MosSCI targeting vector backbone generated as two amplicons from pGH486 (Hollopeter et al., 2014), using primer pairs oGH1011+oEP392 and oGH1012+oEP391, in a three-piece Gibson assembly reaction. The plasmids generated were: pEP29 for *C. elegans* NCAP-1, pEP58 for *M. musculus* NECAP1, pEP41 for *M. musculus* NECAP2, and pEP71 for *S. stellatus* NECAP. These included targeting sequences corresponding to the *cxTi10816* locus and were inserted into the genomes of EG6703 worms as described (Frokjaer-Jensen et al., 2012). The resulting transgenes drive expression of RFP-tagged NECAPs from a ubiquitous *C. elegans* promoter (*Pdpy-30*). Strains, plasmids, and oligonucleotides are listed in Supplementary file 1.

### Tissue culture pulldowns

The heterologous expression of affinity-tagged proteins in HEK293 cells followed by HaloTag isolation from cell lysates and western blots analysis of the purified proteins was performed as described (Hollopeter et al., 2014). Mammalian vectors for the expression of HaloTag fusions with the worm and mouse NECAPs were generated by amplifying the coding sequences from cDNA and inserting them by Gibson assembly into a custom-built HaloTag expression vector (Banks et al., 2014) that was amplified using oGH953-4. The *C*. *elegans* NCAP-1 (NM_061997.5; amplicon oGH955-6) expression vector is pGH500, while the *M*. *musculus* NECAP1 (BK000656.1; amplicon oGH957-8) is pGH501, and *M*. *musculus* NECAP2 (BK000657.1; amplicon oGH959-60) is pGH502. Plasmids and oligonucleotides are listed in Supplementary files 1B and 1C, respectively.

### Recombinant AP2 cores

Bicistronic vectors expressing the hexahistidine-tagged mouse AP2 β2 trunk along with mouse μ2 were based on pGH424 (Hollopeter et al., 2014) and generated as follows: Mutations were introduced in μ2 using Gibson assembly, and primers oGH847-8 for E302K and oGB24+33 for T156A. A thrombin protease site (not utilized in this study) was inserted after amino acid 236 of μ2 using oGB34-35. The construct encoding wild type β2 trunk/μ2 is pGB21, while E302K is pGB19 and T156A is pGB31. The bicistronic vector expressing the GST-tagged mouse AP2 *α* trunk along with rat *σ*2 (pGH504) was generated as follows: The *α* trunk/*σ*2 hemicomplex was amplified from the original expression vector (Collins et al., 2002) using primers oGH1249-50 and recombined with the pACYCDuet vector backbone (amplicon oGH1246-47) using Gibson assembly. The thrombin cleavage site between the *α* trunk and GST was replaced with the human rhinovirus (HRV) C3 protease site (not utilized in this study) with oGH1227-8 and Gibson assembly. Simultaneous expression of all four subunits of the AP2 core was achieved by transforming *E. coli* BL21(DE3) cells (New England Biolabs, Ipswich, MA) with a pair of the bicistronic vectors and co-selecting for ampicillin resistance (pGB21, pGB19 or pGB31) and chloramphenicol resistance (pGH504). To generate phosphorylated AP2 cores, the kinase domain of mouse AAK1 (amino acids 1-325, BC141176.1) was amplified from plasmid template using oEP17-18 and inserted into the pRSFDuet backbone amplified with oEP13 and oGH338 using Gibson assembly. The AP2 subunits were co-expressed with this vector (pEP82) by including selection for kanamycin resistance. Plasmids and oligonucleotides are listed in Supplementary files 1B and 1C, respectively.

2xYT culture media containing the appropriate antibiotics was inoculated with an overnight culture of bacterial cells expressing AP2 cores and incubated at 37°C with shaking at 180-200 RPM until OD_600_ = ∼1.0. The temperature was decreased to 18°C for 1 h before expression was induced with 100 μM isopropyl *β*-D-1-thiogalactopyranoside (IPTG) for 20-24 h. Cells were harvested by centrifugation, washed with lysis buffer (see below), and snap frozen in liquid N2 prior to storage at -80°C. Cell pellets were resuspended in 50 mL lysis buffer per liter of initial culture volume. GST lysis buffer consists of PBS pH 7.4, 1 mM DTT, 60 μg/mL DNase I (grade II from bovine pancreas, Roche), 2.5 mM MgCl_2_, 0.3 mg/mL lysozyme (Sigma), and 1 mM phenylmethylsulfonyl fluoride (PMSF) (Millipore, Billerica, MA) with the addition of 1 tablet of cOmplete EDTA-free Protease Inhibitor Cocktail (Roche) for every 100 mL. The cell slurry was sonicated (20 s pulses at 20% amplitude for a total of 8 min; 50 mL at a time) using a Q700 sonicator (Qsonica, Newtown, CT). Lysates were cleared by centrifugation (∼20000 x g) and filtration (0.2 μm). Cleared filtrate was rotated for 1 h at 4°C with equilibrated GST resin (GE Healthcare Life Sciences, Uppsala, Sweden), 1 ml resin per liter of the initial culture volume. The filtrate and resin were poured over a gravity column, washed with PBS pH 7.4 + 1 mM DTT, and AP2 was eluted with 50 mM Tris pH 8.0 with 10 mM reduced glutathione (Fischer BioReagents, Fairlawn, NJ) and 1 mM DTT. The eluate buffer was exchanged with TBS pH 7.6+ 1 mM DTT and AP2 cores were concentrated to ∼0.5 mg/mL using an Amicon Centrifugal Filter Unit with a 100 kDa cutoff (Merck Millipore). Aliquots of AP2 cores were snap frozen in liquid N2 prior to storage at -80°C.

### Recombinant NECAPs

The NECAP used in Figure 4C had an N-terminal hexahistidine tag, HaloTag and TEV protease site. Mouse NECAP2 (BK000657.1) was amplified from pGH502 using primers oGH1205-6, and recombined with the backbone of the 6xHis-HaloTag-APA expression vector pGH493 (Hollopeter et al., 2014) (amplified with oGH853+1204) using Gibson assembly. This replaced the APA domain with NECAP2 and generated pGH503.

Because we sometimes noted heterogeneity in our recombinant NECAPs, we built a set of vectors to purify NECAPs with cleavable C-terminal affinity tags. The NECAPs used for pulldowns in Figure 4B were purified using the Intein Mediated Purification with an Affinity Chitin-binding Tag (IMPACT, NEB) system, with slight modifications. The *Mxe* GyrA intein tag and chitin binding domain (CBD) from pTXB1 (NEB) were amplified using oGB52-3 (which adds a hexahistidine tag onto the C-terminus of the CBD). This amplicon was inserted into the pET21b expression vector (PCR product of oGB47 and oGH1231) in a three-piece Gibson reaction along with one of the following NECAPs: pGB29 included *C. elegans* NCAP-1 (NM_061997.5, amplified with oGB48-49 from pGH500), pGB27 included *M. musculus* NECAP1 (BK000656.1, amplified with oGB48+50 from pGH501) and pGB28 included *M. musculus* NECAP2 (BK000657.1, amplified with oGB48+51 from pGH502). The NECAPs were amplified such that they include the N-terminal HaloTag and TEV protease site from the mammalian expression vectors (see Tissue culture pulldowns). All plasmids and oligonucleotides are listed in Supplementary file 1B and 1C, respectively.

*E. coli* BL21(DE3) cells (NEB) expressing HaloTag-NECAPs-intein-6xHis (pGB27-9), 6xHis-HaloTag-NECAP2 (pGH503), or the 6xHis-HaloTag control (pGH494) (Hollopeter et al., 2014) were cultured in Terrific Broth (RPI) containing carbenicillin selection. Cultures were subsequently grown as described for AP2 cores (above) except expression was induced with 300 μM IPTG and cells were harvested after 16-20 h incubation at 18°C.

For 6xHis-HaloTag-NECAP2 (pGH503) and 6xHis-HaloTag (pGH494) cell pellets collected from 500 mL of culture re-suspended in 50 mL nickel lysis buffer (20 mM HEPES pH 7.5, 500 mM NaCl, 30 mM imidazole, 5% glycerol, 5 mM BME) with the addition of 60-200 μg/mL DNase I (grade II from bovine pancreas, Roche), 2.5 mM MgCl_2_, 1 tablet of cOmplete EDTA-free Protease Inhibitor Cocktail (Roche), 0.3 mg/mL lysozyme (Sigma), 1 mM PMSF. Cells were lysed by sonication using a Q700 sonicator (Qsonica) at 20% amplitude with 20 sec pulses for 4 min total, centrifuged at ∼20000 x g for 30 min and filtered to 0.2 μm. The hexahistidine-tagged proteins were purified using nickel-charged 5 mL HiTrap Chelating HP columns (GE Healthcare Life Sciences) on a BioLogic LP system (BioRad). After sample loading, columns were washed with nickel lysis buffer and bound proteins were eluted with nickel elution buffer (nickel lysis buffer with 1 M imidazole). Fractions (1 mL each) containing the protein of interest were combined, buffer exchanged with 20 mM Tris pH 8, 5% glycerol, 1 mM EDTA, 1 mM DTT, and 50 mM NaCl and purified by ion exchange chromatography using a HiTrap Q HP column (GE Healthcare Life Sciences) on the BioLogic LP system (BioRad). After sample loading, proteins were eluted using a salt gradient (50 mM to 1000 mM NaCl). Elution fractions containing the protein of interest were combined and aliquots were snap frozen in liquid N2 prior to storage at -80°C.

To purify recombinant NECAPs (HaloTag-NECAPs-intein-6xHis) we used a modified version of the IMPACT system. Cell lysis was performed as described for other hexahistidine-tagged proteins (above) except BME was omitted from the lysis buffer to inhibit intein self-cleavage. Cleared lysates were loaded onto gravity columns containing nickel resin (Thermo Scientific, 0.5 mL per 500 mL initial culture volume) and washed extensively. Columns were then flushed with nickel lysis buffer containing 40 mM BME, plugged, and incubated at 4°C for 30 to 40 h in order to cleave proteins from the intein tag. The released proteins were collected, buffer exchanged with TBS pH 7.6 + 1 mM DTT, and concentrated using an Amicon Centrifugal Filter Unit with 50 kDa MW cutoff (Merck Millipore). Aliquots were snap frozen in liquid N2 prior to storage at -80°C.

### *In vitro* Pulldown Assays

Pulldown assays were performed essentially as described in Hollopeter et al. (2014) except the protease cleavage step was 3 h and the input gel sample represented 20% of the ‘prey’ protein mixture.

### Nerve Ring Microscopy

Worms expressing APA-2:GFP (*mewSi1*) and RFP:NCAP-1 (*mewSi2*) in an *ncap-1* background were imaged on an Zeiss LSM 880 confocal microscope (Biotechnology Resource Center) with a 40x water immersion objective. APA-2:GFP(*mewSi1*) is molecularly similar to APA-2:GFP(*oxSi254*) (Gu et al., 2013) except the entry clones ([4-1]*Pdpy-30*, [1-2]*apa-2* cDNA, and [2-3]*GFP:unc-54 3’UTR*) were recombined with the [4-3] MosSCI vector (pCFJ210) to target the ttTi4348 site in EG6701 worms (Frokjaer-Jensen et al., 2012). Fluorophores were excited with 488 nm (GFP) and 561 nm (RFP) lasers. All strains were imaged in one session with the same laser settings. For each worm, a single confocal slice through the approximate sagittal section of the nerve ring was analyzed in Fiji (Schindelin et al., 2012). Two regions of interest (ROI) corresponding to both dorsal and ventral sections of the nerve ring along with an ROI outside of the worm (to control for background signal) were user defined. The average pixel intensities in both the GFP and RFP channels were determined and the background values were subtracted from the nerve ring values.

### CRISPR-Cas9 generation of μ2 mutations

CRISPR/Cas9 edits were generated using the *dpy-10* co-conversion strategy (Arribere et al., 2014) with ribonucleoprotein (RNP) complexes (Paix, Folkmann, Rasoloson, & Seydoux, 2015). Gonads of young adult hermaphrodites were injected with RNP mixes containing ∼3.7 μg/μL Cas9 (purified in-house), 1 μg/μL tracrRNA, 0.08 μg/μL *dpy-10* crRNA, 16.7 μM *dpy-10(cn64)* roller repair, 0.4 μg/μL gene-specific crRNA, and 16.7 μM gene-specific repair. The gene-specific crRNAs and oligonucleotide repairs were as follows: to generate μ2(E306K) – rEP360 and oGB154 (introduces an *Nru*I site for genotyping), to generate μ2(T160A) – rGB156 and oGB130 (introduces an *Ngo*MIV site), and to generate μ2(R440S) – rGB155 and oGB159 (introduces a *Pvu*I site). F1 progeny exhibiting the roller phenotype were placed on individual culture plates and allowed to produce offspring for 1-2 days before being lysed in 50 μL 1X Phusion GC buffer (NEB) with 0.4 U Proteinase K (NEB) by freezing at -80°C and then heating at 65°C for 1 hr followed by 95°C for 15 min. The targeted region of the genome was amplified using PCR and the amplicon was digested to identify correctly edited worms. Mutations were confirmed by sequencing the PCR product.

## ADDITIONAL FILES

Supplementary file 1. A. Strains B. Plasmids C. Oligonucleotides.

## ACKNOWLEDGEMENTS

We thank Erik Jorgensen, Alejandro Sánchez Alvarado, and the Stowers Institute for Medical Research for support. Ho Yi Mak provided lab space, reagents, and technical advice during the early stages of this work at the Stowers Institute. Technical assistance at the Stowers Institute was provided by Jim Vallandingham (Bioinformatics), Kendra Walton (Molecular Biology), and Valerie Neubauer (Tissue Culture). We thank the labs of Maurine Linder, Joshua Chappie, Carolyn Sevier, Hector Aguilar-Carreno, and Carrie Adler for technical advice and the use of equipment. We thank Anthony Bretscher, Carolyn Sevier, Chris Fromme, Joshua Chappie, Carrie Adler, and Holger Sondermann for constructive criticism that greatly improved the manuscript. G.M.B. and E.A.P. were supported by an NIH training grant GM007273-43. G.M.B. is supported by an NSF graduate research fellowship DGE-1650441.

---

**Figure 1–figure supplement 1.**
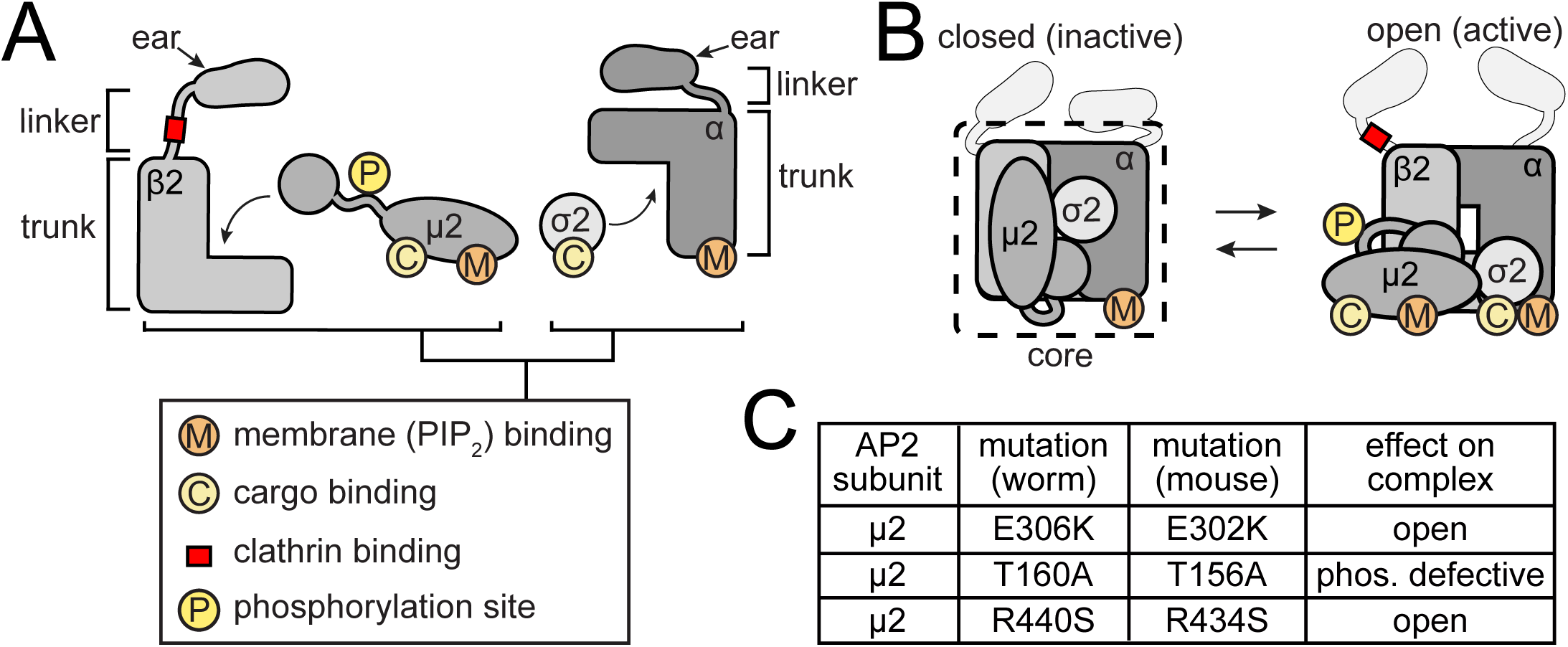
AP2 structures and mutations. **A.** AP2 is comprised of two large adaptins (α and β2) and two smaller subunits (μ2 and σ2). The adaptins, in turn, are comprised of appendage (ear), hinge (linker) and trunk domains. Phosphorylation site and binding pockets are diagrammed. **B.** Cartoon representations of the closed (Collins *et al*., 2002) and open (Jackson *et al*., 2010 and Kelly *et al*., 2014) AP2 conformations. The core complex (dashed line) lacks ears and linkers. **C.** Table of AP2 mutations used in this study.

## REFERENCES

Banks, C. A., Lee, Z. T., Boanca, G., Lakshminarasimhan, M., Groppe, B. D., Wen, Z., Hattem, G. L., Seidel, C. W., Florens, L., & Washburn, M. P. (2014). Controlling for gene expression changes in transcription factor protein networks. Mol Cell Proteomics, 13(6), 1510–1522. doi:10.1074/mcp.M113.033902

Brenner, S. (1974). The genetics of Caenorhabditis elegans. Genetics, 77(1), 71–94.

Chamberland, J. P., Antonow, L. T., Dias Santos, M., & Ritter, B. (2016). NECAP2 controls clathrin coat recruitment to early endosomes for fast endocytic recycling. J Cell Sci, 129(13), 2625–2637. doi:10.1242/jcs.173708

Chappell, T. G., Welch, W. J., Schlossman, D. M., Palter, K. B., Schlesinger, M. J., & Rothman, J. E. (1986). Uncoating ATPase is a member of the 70 kilodalton family of stress proteins. Cell, 45(1), 3–13.

Cocucci, E., Aguet, F., Boulant, S., & Kirchhausen, T. (2012). The first five seconds in the life of a clathrin-coated pit. Cell, 150(3), 495–507. doi:10.1016/j.cell.2012.05.047

Collins, B. M., McCoy, A. J., Kent, H. M., Evans, P. R., & Owen, D. J. (2002). Molecular architecture and functional model of the endocytic AP2 complex. Cell, 109(4), 523–535.

Conner, S. D., & Schmid, S. L. (2002). Identification of an adaptor-associated kinase, AAK1, as a regulator of clathrin-mediated endocytosis. J Cell Biol, 156(5), 921–929. doi:10.1083/jcb.200108123

Conner, S. D., Schroter, T., & Schmid, S. L. (2003). AAK1-mediated micro2 phosphorylation is stimulated by assembled clathrin. Traffic, 4(12), 885–890.

Cremona, O., Di Paolo, G., Wenk, M. R., Luthi, A., Kim, W. T., Takei, K., Daniell, L., Nemoto, Y., Shears, S. B., Flavell, R. A., McCormick, D. A., & De Camilli, P. (1999). Essential role of phosphoinositide metabolism in synaptic vesicle recycling. Cell, 99(2), 179–188.

Davis, M. W., Hammarlund, M., Harrach, T., Hullett, P., Olsen, S., & Jorgensen, E. M. (2005). Rapid single nucleotide polymorphism mapping in C. elegans. BMC Genomics, 6, 118. doi:10.1186/1471-2164-6-118

Dergai, M., Iershov, A., Novokhatska, O., Pankivskyi, S., & Rynditch, A. (2016). Evolutionary Changes on the Way to Clathrin-Mediated Endocytosis in Animals. Genome Biol Evol, 8(3), 588–606. doi:10.1093/gbe/evw028

Ehrlich, M., Boll, W., Van Oijen, A., Hariharan, R., Chandran, K., Nibert, M. L., & Kirchhausen, T. (2004). Endocytosis by random initiation and stabilization of clathrin-coated pits. Cell, 118(5), 591–605. doi:10.1016/j.cell.2004.08.017

Frokjaer-Jensen, C., Davis, M. W., Ailion, M., & Jorgensen, E. M. (2012). Improved Mos1-mediated transgenesis in C. elegans. Nat Methods, 9(2), 117–118. doi:10.1038/nmeth.1865

Frokjaer-Jensen, C., Davis, M. W., Hopkins, C. E., Newman, B. J., Thummel, J. M., Olesen, S. P., Grunnet, M., & Jorgensen, E. M. (2008). Single-copy insertion of transgenes in Caenorhabditis elegans. Nat Genet, 40(11), 1375–1383. doi:10.1038/ng.248

Ghosh, P., & Kornfeld, S. (2003). AP-1 binding to sorting signals and release from clathrin-coated vesicles is regulated by phosphorylation. J Cell Biol, 160(5), 699–708. doi:10.1083/jcb.200211080

Gibson, D. G., Young, L., Chuang, R. Y., Venter, J. C., Hutchison, C. A., 3rd, & Smith, H. O. (2009). Enzymatic assembly of DNA molecules up to several hundred kilobases. Nat Methods, 6(5), 343–345. doi:10.1038/nmeth.1318

Greener, T., Zhao, X., Nojima, H., Eisenberg, E., & Greene, L. E. (2000). Role of cyclin G-associated kinase in uncoating clathrin-coated vesicles from non-neuronal cells. J Biol Chem, 275(2), 1365–1370.

Gu, M., Liu, Q., Watanabe, S., Sun, L., Hollopeter, G., Grant, B. D., & Jorgensen, E. M. (2013). AP2 hemicomplexes contribute independently to synaptic vesicle endocytosis. Elife, 2, e00190. doi:10.7554/eLife.00190

Hannan, L. A., Newmyer, S. L., & Schmid, S. L. (1998). ATP- and cytosol-dependent release of adaptor proteins from clathrin-coated vesicles: A dual role for Hsc70. Mol Biol Cell, 9(8), 2217–2229.

Henne, W. M., Boucrot, E., Meinecke, M., Evergren, E., Vallis, Y., Mittal, R., & McMahon, H. T. (2010). FCHo proteins are nucleators of clathrin-mediated endocytosis. Science, 328(5983), 1281–1284. doi:10.1126/science.1188462

Hollopeter, G., Lange, J. J., Zhang, Y., Vu, T. N., Gu, M., Ailion, M., Lambie, E. J., Slaughter, B. D., Unruh, J. R., Florens, L., & Jorgensen, E. M. (2014). The membrane-associated proteins FCHo and SGIP are allosteric activators of the AP2 clathrin adaptor complex. Elife, 3. doi:10.7554/eLife.03648

Honing, S., Ricotta, D., Krauss, M., Spate, K., Spolaore, B., Motley, A., Robinson, M., Robinson, C., Haucke, V., & Owen, D. J. (2005). Phosphatidylinositol-(4,5)-bisphosphate regulates sorting signal recognition by the clathrin-associated adaptor complex AP2. Mol Cell, 18(5), 519–531. doi:10.1016/j.molcel.2005.04.019

Jackson, A. P., Flett, A., Smythe, C., Hufton, L., Wettey, F. R., & Smythe, E. (2003). Clathrin promotes incorporation of cargo into coated pits by activation of the AP2 adaptor micro2 kinase. J Cell Biol, 163(2), 231–236. doi:10.1083/jcb.200304079

Jackson, L. P., Kelly, B. T., McCoy, A. J., Gaffry, T., James, L. C., Collins, B. M., Honing, S., Evans, P. R., & Owen, D. J. (2010). A large-scale conformational change couples membrane recruitment to cargo binding in the AP2 clathrin adaptor complex. Cell, 141(7), 1220– 1229. doi:10.1016/j.cell.2010.05.006

Kadlecova, Z., Spielman, S. J., Loerke, D., Mohanakrishnan, A., Reed, D. K., & Schmid, S. L. (2017). Regulation of clathrin-mediated endocytosis by hierarchical allosteric activation of AP2. J Cell Biol, 216(1), 167–179. doi:10.1083/jcb.201608071

Kelly, B. T., Graham, S. C., Liska, N., Dannhauser, P. N., Honing, S., Ungewickell, E. J., & Owen, D.J. (2014). Clathrin adaptors. AP2 controls clathrin polymerization with a membrane-activated switch. Science, 345(6195), 459–463. doi:10.1126/science.1254836

Kelly, B. T., McCoy, A. J., Spate, K., Miller, S. E., Evans, P. R., Honing, S., & Owen, D. J. (2008). A structural explanation for the binding of endocytic dileucine motifs by the AP2 complex. Nature, 456(7224), 976–979. doi:10.1038/nature07422

Kim, E., Sun, L., Gabel, C. V., & Fang-Yen, C. (2013). Long-term imaging of Caenorhabditis elegans using nanoparticle-mediated immobilization. PLoS One, 8(1), e53419. doi:10.1371/journal.pone.0053419

Kirchhausen, T., Owen, D., & Harrison, S. C. (2014). Molecular structure, function, and dynamics of clathrin-mediated membrane traffic. Cold Spring Harb Perspect Biol, 6(5), a016725. doi:10.1101/cshperspect.a016725

Manna, P. T., Gadelha, C., Puttick, A. E., & Field, M. C. (2015). ENTH and ANTH domain proteins participate in AP2-independent clathrin-mediated endocytosis. J Cell Sci, 128(11), 2130– 2142. doi:10.1242/jcs.167726

Matsui, W., & Kirchhausen, T. (1990). Stabilization of clathrin coats by the core of the clathrin-associated protein complex AP-2. Biochemistry, 29(48), 10791–10798.

Olusanya, O., Andrews, P. D., Swedlow, J. R., & Smythe, E. (2001). Phosphorylation of threonine 156 of the mu2 subunit of the AP2 complex is essential for endocytosis in vitro and in vivo. Curr Biol, 11(11), 896–900.

Paix, A., Folkmann, A., Rasoloson, D., & Seydoux, G. (2015). High Efficiency, Homology-Directed Genome Editing in Caenorhabditis elegans Using CRISPR-Cas9 Ribonucleoprotein Complexes. Genetics, 201(1), 47–54. doi:10.1534/genetics.115.179382

Rapoport, I., Miyazaki, M., Boll, W., Duckworth, B., Cantley, L. C., Shoelson, S., & Kirchhausen, T. (1997). Regulatory interactions in the recognition of endocytic sorting signals by AP-2 complexes. EMBO J, 16(9), 2240–2250. doi:10.1093/emboj/16.9.2240

Reider, A., Barker, S. L., Mishra, S. K., Im, Y. J., Maldonado-Baez, L., Hurley, J. H., Traub, L. M., & Wendland, B. (2009). Syp1 is a conserved endocytic adaptor that contains domains involved in cargo selection and membrane tubulation. EMBO J, 28(20), 3103–3116. doi:10.1038/emboj.2009.248

Ricotta, D., Conner, S. D., Schmid, S. L., von Figura, K., & Honing, S. (2002). Phosphorylation of the AP2 mu subunit by AAK1 mediates high affinity binding to membrane protein sorting signals. J Cell Biol, 156(5), 791–795. doi:10.1083/jcb.200111068

Ritter, B., Denisov, A. Y., Philie, J., Deprez, C., Tung, E. C., Gehring, K., & McPherson, P. S. (2004). Two WWF-based motifs in NECAPs define the specificity of accessory protein binding to AP-1 and AP-2. EMBO J, 23(19), 3701–3710. doi:10.1038/

Ritter, B., Murphy, S., Dokainish, H., Girard, M., Gudheti, M. V., Kozlov, G., Halin, M., Philie, J., Jorgensen, E. M., Gehring, K., & McPherson, P. S. (2013). NECAP 1 regulates AP-2 interactions to control vesicle size, number, and cargo during clathrin-mediated endocytosis. PLoS Biol, 11(10), e1001670. doi:10.1371/journal.pbio.1001670

Ritter, B., Philie, J., Girard, M., Tung, E. C., Blondeau, F., & McPherson, P. S. (2003). Identification of a family of endocytic proteins that define a new α-adaptin ear-binding motif. EMBO reports, 4(11), 1089–1093. doi:10.1038/sj.embor.7400004

Sakaushi, S., Inoue, K., Zushi, H., Senda-Murata, K., Fukada, T., Oka, S., & Sugimoto, K. (2007). Dynamic behavior of FCHO1 revealed by live-cell imaging microscopy: its possible involvement in clathrin-coated vesicle formation. Biosci Biotechnol Biochem, 71(7), 1764–1768. doi:10.1271/bbb.60720

Sato, K., Norris, A., Sato, M., & Grant, B. D. (2014). C. elegans as a model for membrane traffic. WormBook, 1–47. doi:10.1895/wormbook.1.77.2

Schindelin, J., Arganda-Carreras, I., Frise, E., Kaynig, V., Longair, M., Pietzsch, T., Preibisch, S., Rueden, C., Saalfeld, S., Schmid, B., Tinevez, J. Y., White, D. J., Hartenstein, V., Eliceiri, K.,Tomancak, P., & Cardona, A. (2012). Fiji: an open-source platform for biological-image analysis. Nat Methods, 9(7), 676–682. doi:10.1038/nmeth.2019

Semerdjieva, S., Shortt, B., Maxwell, E., Singh, S., Fonarev, P., Hansen, J., Schiavo, G., Grant, B. D., & Smythe, E. (2008). Coordinated regulation of AP2 uncoating from clathrin-coated vesicles by rab5 and hRME-6. J Cell Biol, 183(3), 499–511. doi:10.1083/jcb.200806016

Taylor, M. J., Perrais, D., & Merrifield, C. J. (2011). A high precision survey of the molecular dynamics of mammalian clathrin-mediated endocytosis. PLoS Biol, 9(3), e1000604. doi:10.1371/journal.pbio.1000604

Uezu, A., Horiuchi, A., Kanda, K., Kikuchi, N., Umeda, K., Tsujita, K., Suetsugu, S., Araki, N., Yamamoto, H., Takenawa, T., & Nakanishi, H. (2007). SGIP1alpha is an endocytic protein that directly interacts with phospholipids and Eps15. J Biol Chem, 282(36), 26481–26489. doi:10.1074/jbc.M703815200

Umasankar, P. K., Ma, L., Thieman, J. R., Jha, A., Doray, B., Watkins, S. C., & Traub, L. M. (2014). A clathrin coat assembly role for the muniscin protein central linker revealed by TALEN-mediated gene editing. Elife, 3. doi:10.7554/eLife.04137

Umeda, A., Meyerholz, A., & Ungewickell, E. (2000). Identification of the universal cofactor (auxilin 2) in clathrin coat dissociation. Eur J Cell Biol, 79(5), 336–342. doi:10.1078/S0171-9335(04)70037-0

Ungewickell, E., Ungewickell, H., Holstein, S. E., Lindner, R., Prasad, K., Barouch, W., Martin, B., Greene, L. E., & Eisenberg, E. (1995). Role of auxilin in uncoating clathrin-coated vesicles. Nature, 378(6557), 632–635. doi:10.1038/378632a0

White, J. G., Southgate, E., Thomson, J. N., & Brenner, S. (1986). The structure of the nervous system of the nematode Caenorhabditis elegans. Philos Trans R Soc Lond B Biol Sci, 314(1165), 1–340.

